# Intrinsic variability of fluorescence calibrators impacts the assignment of MESF or ERF values to nanoparticles and extracellular vesicles by flow cytometry

**DOI:** 10.1101/2021.03.01.433358

**Authors:** Estefanía Lozano-Andrés, Tina Van Den Broeck, Lili Wang, Majid Mehrpouyan, Ye Tian, Xiaomei Yan, Marca H.M. Wauben, Ger. J.A. Arkesteijn

## Abstract

Flow cytometry is commonly used to characterize nanoparticles (NPs) and extracellular vesicles (EVs) but results are often expressed in arbitrary units to indicate fluorescence intensity. This hampers interlaboratory and inter-platform comparisons. We investigated the use of molecules of equivalent soluble fluorophores (MESF)-beads for assignment of fluorescence values to NPs and EVs by comparing two FITC-MESF bead sets as calibrators on different flow cytometry platforms (BD Influx^™^, CytoFLEX LX^™^ and SORP BD FACSCelesta^™^). Next, fluorescence signals of NPs and EVs were calibrated using different sets of FITC and PE-MESF beads. Fluorescence calibration using beads designed for cellular flow cytometry allowed inter-platform comparison. However, the intrinsic uncertainty in the fluorescence assignment to these MESF beads impacts the reliable assignment of MESF values to NPs and EVs based on extrapolation into the dim fluorescence range. Our findings demonstrate that the use of the same set of calibration materials (vendor and lot number) and the same number of calibration points, greatly improves robust interlaboratory and inter-platform comparison of fluorescent submicron sized particles.

## Introduction

A well-known fluorescence calibration method in flow cytometry (FC) is the use of fluorescent beads to which a measurement value is assigned using standardized units established by the National Institute of Standards and Technology (NIST), such as molecules of equivalent soluble fluorophores (MESF) or equivalent number of reference fluorophore (ERF). This calibration method was developed for cellular FC and allows for quantifiable fluorescence measurements and platform comparison. The fluorescence intensity on the calibrator beads matches the expected intensity on the labeled cells. Therefore, the calibrated cellular fluorescence variation closely compares to the intrinsic variation on the calibrator and allows for data interpolation [1–4]. In 2012, a NIST/ISAC standardization study reported differences in the assigned units to calibrators from different manufactures, indicating the importance of the examination of the accuracy and precision of available calibrators [5]. Nevertheless, the assignment of a specific MESF or ERF value to calibration beads is inextricably bound to a variation around this value. This variation translates in an uncertainty level between the measured and the assigned values that remains acceptable as long as the sample values are within the range of the calibrator.

During the last decade, small particle FC has become a powerful tool for high-throughput analysis of nanoparticles (NPs) and cell-derived extracellular vesicles (EVs) [6, 7]. However, EV measurements are challenging, mainly because the vast majority of EVs is small in size (<200nm) and their light scattering and fluorescent signals are typically close to, at, or below the instrument’s detection limit [8]. Furthermore, the majority of data is reported in arbitrary units of fluorescence, which is cumbersome for the analysis of dim and small particles, whereby particles cannot be fully discriminated from negative counterparts and background signals. The MIFlowCyt-EV framework recommends the use of MESF beads for calibration and standardized reporting of EV flow cytometric experiments, especially when a fluorescent threshold is applied [8]. However, since available calibrators are developed for cells and as such are much brighter in fluorescence than EVs it is unknown to which extend these calibrators will provide precision and/or accuracy for the assignment of fluorescent values to NPs and EVs. We investigated how the given units of the calibrator impact the regression line for assignment of MESF and/or ERF units to NPs and EVs. Therefore, we evaluated custom-made calibrator beads sets from the same manufacturer on three different flow cytometers and provide insights on how different bead sets affect the calibration of fluorescence signals from NPs and EVs.

## Results

### Assessment of precision and accuracy of different MESF bead sets for fluorescence calibration across platforms

To assess the precision and accuracy of MESF bead sets for fluorescence calibration across platforms, two FITC MESF bead sets of 6 μm and 2 μm, containing respectively five or four fluorescent bead populations, were selected for measurements on three different instruments, namely a BD Influx, a BC CytoFLEX and a SORP BD FACSCelesta ^™^. Since calibrator bead sets can differ in the number of fluorescent bead populations (typically ranging from 3 to 5) and the number of calibrator points can impact the slope of the regression line (Figure S1a-b), we included for fluorescence calibration across platforms equal numbers of fluorescent bead populations (n=4) of the two FITC MESF bead sets that were consistently measured on all three platforms (Figure 1).

**Figure 1.**
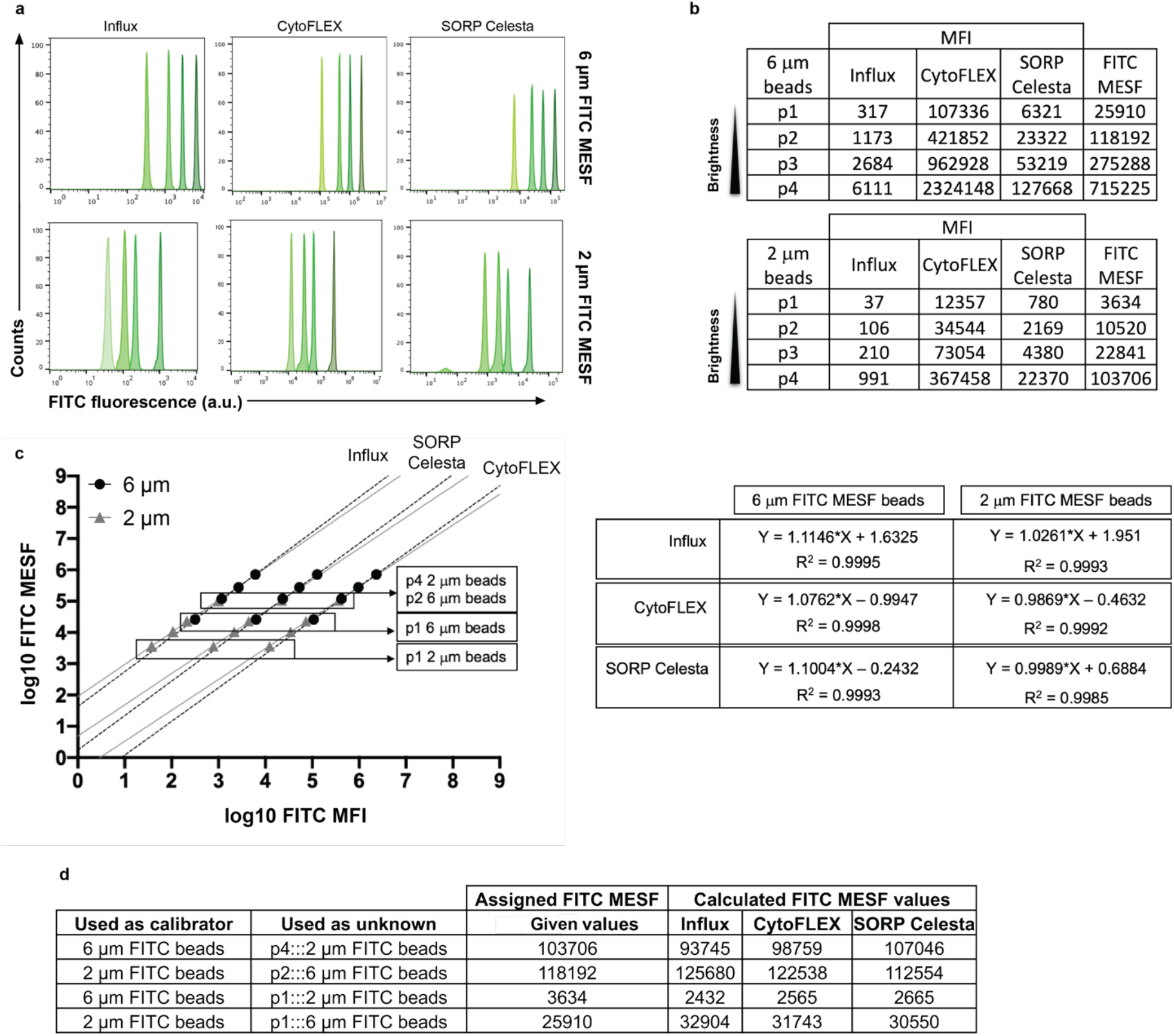
Evaluation of two different FITC MESF bead sets for the calibration of fluorescent intensities across three flow cytometer platforms. **(a)** Histogram overlays (axis in arbitrary units) of FITC fluorescent intensity peaks derived from the 6 μm (upper row) or the 2 μm (lower row) FITC MESF beads **(b)** Table showing the median fluorescence intensity (MFI) statistic derived from each of the fluorescent intensity peaks from dimmer to brighter being expressed in arbitrary units as well as the assigned MESF values (right column). **(c)** Least square linear regression analysis of 6 μm (black circles) and 2 μm (grey triangles) FITC MESF beads. Provided FITC MESF and measured FITC MFI values were transformed to log and plotted in a log-log fashion for the three platforms. **(d)** Table indicating the expected and calculated FITC MESF values for each sample used in the analysis.

Singlet gated populations are displayed as overlays in histograms showing the FITC fluorescence (Figure 1a) and indicated from dim to bright as p1, p2, p3 and p4 (Figure 1b). The 6 μm FITC MESF beads contained overall brighter fluorescent intensities, whose assigned values range from 25,910 to 715,225 FITC MESF, while the 2 μm FITC MESF beads covered a dimmer part of the fluorescence intensity range with assigned values ranging from 3,634 to 103,706 FITC MESF. Clearly, MFI arbitrary units cannot be directly compared between instruments (Figure 1b), but after fluorescence calibration comparable FITC MESF units could be assigned (Figure 1c-d). Nevertheless, the two calibration bead sets displayed a different slope with a consistent tendency across the three platforms, suggesting a variation introduced by an inherent attribute of the beads themselves. This lead us to further examine the robustness of the calibration. To gain insight into the precision and accuracy of MESF assignments we selected specific bead populations from one set, referred to as ‘unknown’ in Figure 1d, to recalculate their FITC MESF units using the regression line from the other bead set. The selected ‘unknown’ samples used were: (i) p1 and p4 of the 2 μm bead set for which the FITC MESF values were calculated using the regression line of the 6 μm bead set and (ii) p1 and p2 of the 6 μm bead set for which the FITC MESF values were calculated using the regression line of the 2 μm bead set (Figure 1c). Using this approach, the calculated MESF values of p4 from the 2 μm beads and p2 from the 6 μm beads showed less than 20% variation (10% above or 10% below actual values) of the actual value, while data were precise when compared between platforms (Figure 1d). Also the MESF values of the dimmest p1 2 μm and 6 μm beads, calculated using respectively the regression lines of the 6 μm and 2 μm bead sets, were comparable between platforms. However, the calculated values revealed more than a 20% variation from the given value, leading to either an underestimation or an overestimation of the FITC MESF units (Figure 1d). These results show that slight differences in the slope of the calibration lines of the different MESF bead sets become more prominent when extrapolation needs to be extended into the dim area beyond the fluorescence intensities of the calibration beads themselves. Since the same slope differences occurred on all three platforms (Figure 1c), this observation is not related to the type of instrument used (e.g.; digital or analog, photomultiplier (PMT) or avalanche photodiode (APD), jet-in-air or cuvette based). Furthermore, this recurring pattern on all platforms makes it unlikely that differences in slopes were caused by instrument non-linearity. Further evidence to rule out non-linearity issues is provided by linear plotting of the values, showing no non-linearity issues (Figure S2a-c), and demonstrating instrument linearity on the BD Influx following the approach described by Bagwell et al [9] (Figure S3). Furthermore, we ruled out that slope differences were a result of variations between separate measurements (Figure S4), and confirmed by testing both custom-made and commercial FITC MESF beads (Figure S1) and FITC MESF beads and PE MESF beads (Figure S2 d-f) that slope variability is inherent to the use of calibrator beads.

### MESF assignments of FITC fluorescence intensities to synthetic silica nanoparticles depend on the MESF-bead calibrator set

We next investigated how calibration with the two FITC MESF bead sets of 6 μm and 2 μm impacts fluorescent assignment to dim fluorescent nanoparticles. For this purpose 550 nm silica NPs containing 6 populations with FITC fluorescence intensities below or within the range of the calibration beads were measured on the BD Influx. Singlets were gated (Figure S5) and an histogram overlay was generated showing 6 different FITC fluorescence intensities (Figure 2a, left). Histograms showing the calibrated FITC MESF units of these silica NPs based on fluorescence calibration with either the 6 μm or 2 μm FITC MESF calibrator bead set are displayed in Figure 2a (respectively middle or right). The obtained MFI and CV values for each silica NP population, as well as the calculated FITC MESF values based on the two calibrator sets are shown in Figure 2b. The calculated FITC MESF values for the silica NPs appeared consistently lower when the regression line of the 6 μm calibration bead sets was used (Figure 2b). This phenomenon is not limited to the use of FITC MESF beads and can solely be explained by the difference in the slope of the regression line of the two calibrator bead sets, as was confirmed by calculating the fluorescent intensity in terms of PE ERF for 200 nm broad spectrum fluorescent polystyrene NPs based on the 6 μm and 2 μm PE MESF calibrator bead sets (Figure S6a). Importantly, multi-intensity peak analysis of the silica NPs revealed that the difference in FITC MESF values of these NPs obtained by the two calibrator bead sets increased in the dimmer range of the fluorescence, with 27.3% variation for the brightest fluorescent peak (p6) to 76.5% variation for the dimmest population of these NPs (Figure 2b). Also the PE ERF values calculated for the relatively dim 200 nm broad spectrum fluorescent polystyrene NPs based on the 6 μm or 2 μm PE MESF calibrator bead sets showed a variation of 41.3% (Figure S6a). These results can be explained by the fact that the differences in calculated values increase by extrapolation into the dim area as a consequence of the differences in the slopes of the regression lines between the calibrator bead sets.

**Figure 2.**
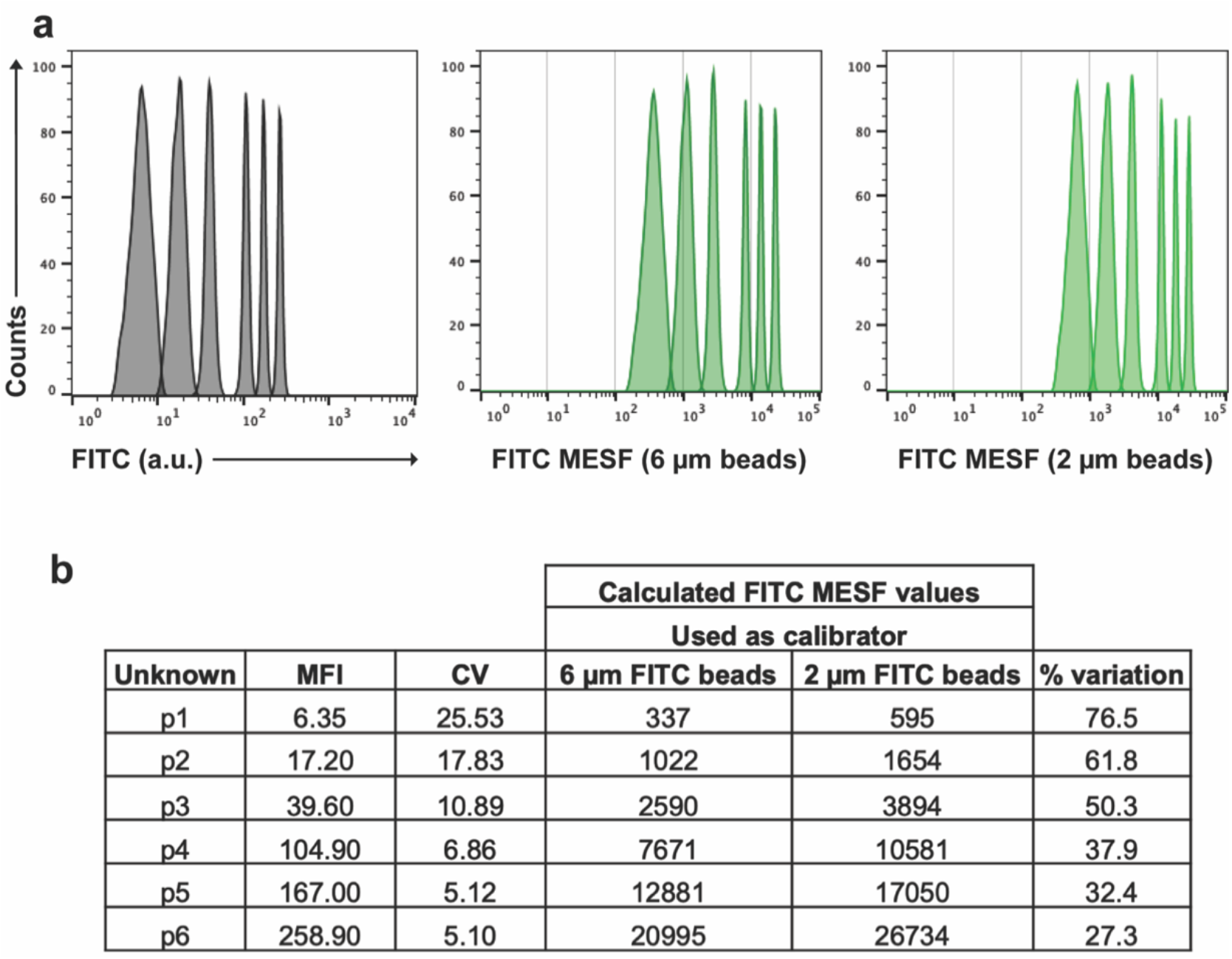
MESF bead-based calibration of fluorescence signals from synthetic silica NPs. **(a)** Histogram overlay showing FITC fluorescence in arbitrary units (a.u.) (left), FITC MESF calibrated axis based on the 6 μm (middle) and 2 μm FITC MESF beads (right) from the six differently FITC-labeled 550 nm NP gated populations. **(b)** Table showing the MFI and CV as well as the calculated FITC MESF values for each of the unknown populations with the percentage of variation between the two calculated reference values.

### MESF calibration using different bead sets leads to variable ERF and MESF values assigned to fluorescently CFSE stained and CD9 labeled extracellular vesicles

We next demonstrate the impact of the assignment of FITC ERF units and PE MESF units to a biological EV sample measured on BD Influx by using the four different MESF calibrator sets, i.e. 6 and 2 μm FITC MESF and the 6 and 2 μm PE MESF beads. Since the light scatter of these EVs was too low to resolve the EV population from the background signals, fluorescence thresholding was applied [10] based on the CFSE luminal dye staining. Furthermore, the expression of CD9, a tetraspanin enriched on the surface of the 4T1-derived EVs, was analyzed by using a CD9-PE antibody (Figure 3a). Unstained EVs and CFSE stained EVs with a matching isotype-PE control were measured side-by-side (Figure 3a) and fluorescent polystyrene spike-in beads (200 nm) were added to EV samples to determine the EV-concentration and define the EV-gating (Figure S7a). In Figure 3c-d, the histogram overlays show how the fluorescence intensities of the calibrators relate to the fluorescent signals generated by CFSE stained and CD9-PE labeled EVs. Our calibration results revealed a 76.6% variation in the calculated CFSE ERF units on CFSE stained EV and a 156.9% variation in the calculated PE MESF units on CD9-PE labeled EVs when the different MESF calibrator sets were used (Figure 3a-b). Moreover, the fluorescent threshold value of 0.67 used on the BD Influx corresponds to an equivalent of 150 FITC MESF based on the 6 μm beads or 300 FITC MESF based on the 2 μm beads (Figure 3a), which also shows the variation between two bead sets when reporting the level of detection in standardized units.

**Figure 3.**
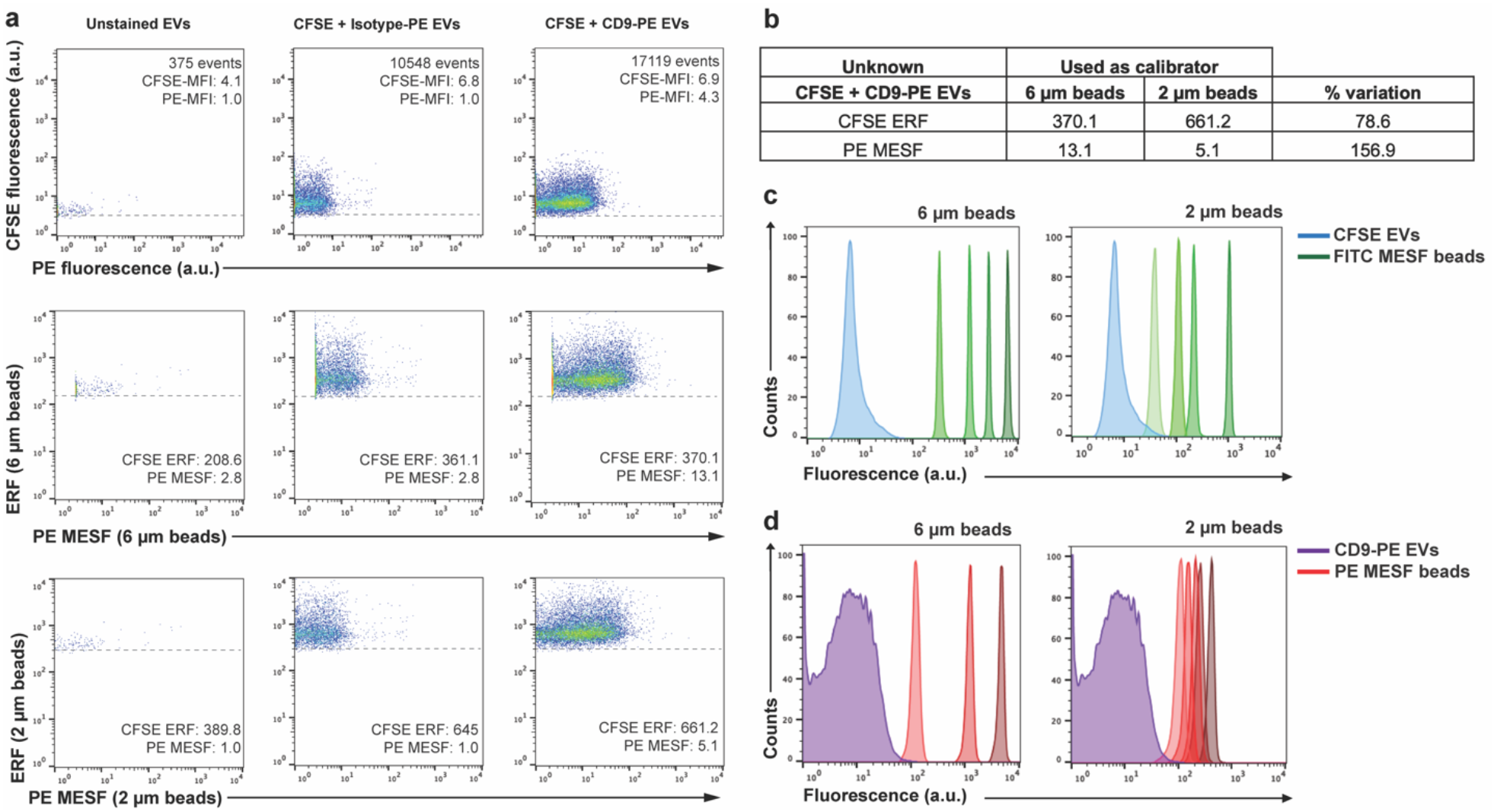
MESF bead-based calibration of fluorescent signals from biological EV samples. **(a)** Analysis of EV samples by using a fluorescence threshold. Unstained EVs control (left), CFSE and isotype-PE stained EVs (middle) and CFSE and CD9-PE stained EVs (right) dot plots showing CFSE fluorescence Vs PE fluorescence in arbitrary units (a.u.) (upper row) or CFSE ERF Vs PE MESF calibrated axis based on either the 6 μm (middle row) or 2 μm calibration beads (lower row). The dashed line in each dot plot indicates the fluorescence threshold value used. Number of events within the EV gate and MFI values for either CFSE or PE fluorescence (a.u.) are indicated in the top row. CFSE ERF and PE MESF values are indicated based on the 6 μm (middle row) or 2 μm beads (lower row). **(b)** Table showing the ERF or MESF values obtained after calibration for the CFSE and CD9-PE stained EVs and the percentage of variation between the use of the 6 μm or 2 μm bead sets. **(c)** Histogram overlays displaying fluorescence in arbitrary units from CFSE stained EVs (blue) next to the 6 μm or 2 μm FITC-MESF bead set (green). **(d)** Histogram overlays displaying fluorescence in arbitrary units from CD9-PE labeled EVs (purple) next to the 6 μm or 2 μm PE-MESF bead set (red).

## Discussion

The field of small particle flow cytometry is rapidly evolving, where the definition of what and how much can be detected is crucial. Besides the inter-comparability of data, the use of MESF/ERF values generates awareness about the range of fluorescence intensities that can be expected for small particles, such as EVs, and allows to indicate instrument detection sensitivity and to report fluorescence thresholding in calibrated units [8, 11].

In line with previous findings, we here showed that linear regression curves derived from calibration beads developed for calibration of fluorescence on cells, can be used to calculate MESF/ERF values for dim NPs and EVs, allowing data inter-comparability with acceptable precision when the same calibrator is used [11]. Since earlier reports pointed out towards variabilities in MESF/ERF assignments between manufacturer’s [5], we used different calibrator bead sets from the same manufacturer, assigned by using the same method and internal NIST traceable calibrator and prepared by following the same manufacturing process. Nevertheless, we found that the robustness of a calculated MESF/ERF value varies upon the use of different calibrator bead sets.

We here demonstrate that the calibration of dim NPs and EVs is substantially affected by the intrinsic variation within the assignment of MESF/ERF values to the calibrator beads [12]. Since the fluorescent intensities of EVs, based on generic staining and/or on antibody labeling, are far dimmer than the available calibrators calculation of their MESF/ERF values relies on extrapolation of the regression line of the calibrator beads into the dim area. Our data demonstrate that regression lines of different calibrator sets result in different calculated MESF/ERF values for NPs and EVs that have fluorescent intensities at the lower end or below the intensities of the calibrator beads themselves. Due to the increased separation of regression lines with slightly different slopes at the lower end, increasing uncertainties exist in the MESF/ERF assignment for dim NPs and EVs, which compromise the accuracy of the MESF/ERF assignment. Importantly, most available calibrator bead sets do not provide uncertainty values around the given MESF/ERF units, which would help to create awareness about the possibilities and limitations related to MESF/ERF unit reporting. Based on our findings, the use of the same calibrator bead set and the same number of data points of the calibrators used for linear regression would increase robustness of the calculation of MESF/ERF values for inter-laboratory and inter-platform comparison, and detailed description of calibration materials and calculation of MESF/ERF values would increase reproducibility.

Clearly, a calibrator with MESF/ERF values closer to the range of fluorescence intensities of the sample of interest and with a low uncertainty of assignment is preferable. Importantly, novel state-of-the-art flow cytometers that are designed to measure small particles rely obligately on sub-micron sized beads to perform MESF/ERF calibration. These state-of-the-art platforms cannot measure the ‘standard’ 6 μm MESF beads [13]. Therefore, there is an urgent need for calibration beads that are validated for small particle flow cytometry.

In summary, our results confirm that fluorescence calibration enables data comparison and provides information on the detection sensitivity of the instrument in standardized units [14, 15], but also urge for awareness of the limitations when fluorescence calibration is being employed for EVs and NPs, especially in terms of accuracy. Lastly, for robust assignments of fluorescence values to NPs and EVs, there is a need for multi-institutional collaborations (between research labs, companies and metrology institutions, such as NIST) to produce and validate calibration materials that have low and well-characterized uncertainty of assigned fluorescent values, ideally allowing for data interpolation and with a size range that is compatible for all flow cytometer platforms.

## Material and Methods

### Calibration beads

For calibration of the fluorescence axis in the fluorescein isothiocyanate (FITC) channel we used two different sets of FITC MESF beads (custom-made, 6 μm lot MM2307 #131-10; #131-8; #130-6; #130-5; #130-3 and 2 μm lot MM2307#156; #159.1; #159.2; #122.3, BD Biosciences, San Jose, CA). For calibration of the fluorescence axis in the PE channel, we used two different sets of PE MESF beads (6 μm, commercial QuantiBrite, Catalog No. 340495 lot 62981 and 2 μm, custom made, lot MM2327#153.1; #153.2; #153.4; #153.5; #153.6, BD Biosciences, San Jose, CA).

The 6 μm FITC MESF beads were prepared by reacting various concentration of FITC with PMMA Beads (Bangs Labs) in borate buffer at pH 9.2. The 2 μm FITC beads were prepared by reacting various concentrations of FITC-BSA (with a FITC/BSA molar ratio of 2) with 2 μm carboxylic beads (Bangs Labs) using EDC/NHS chemistry. The 2 μm PE beads were made as described above except various concentrations of PE were used with 2 μm carboxylic beads in EDC/NHS chemistry. These beads were analyzed on a BD LSRFortessa^™^ (BD Biosciences) and their MESF values were assigned by cross-calibration using commercially available MESF beads (Flow Cytometry Standards Corp.). The FITC ERF values were assigned to both 2 μm and 6 μm beads using a specific lot of FITC-FC Bead (BD Biosciences) as a calibrator with known ERF value, which has been assigned by NIST. This provided us with two distinct calibrator bead sets that were produced through the same manufacturing process and assigned using the same instruments and the same internal NIST traceable calibrator to exclude internal processing variations.

In addition, we measured commercially available Quantum^™^ FITC-5 MESF (7 μm, Catalog No. 555, lot 14609, Bangs Laboratories) and AccuCheck ERF Reference Particles Kit (3 μm, Catalog No. A55950, lot #081220207, #081220203, #081220208, Thermo Fisher) which were prepared according to the manufacturer’s instructions.

All calibration bead sets were measured with gain or voltage settings as would be used for the analysis of small particles (i.e. EVs). In addition, not all beads could be measured on every instrument. For fair cross-platform comparison of the slopes of the regression lines, only the bead populations that could be measured on all instruments were included for linear regression analysis.

### Flow cytometer platforms

In this study three flow cytometers were used. A jet in air-based BD Influx (BD Biosciences, San Jose, CA), a BC CytoFLEX LX (Beckman Coulter, Brea, CA) with a cuvette-based system and a cuvette-based SORP BD FACSCelesta^™^ (BD Biosciences, San Jose, CA) equipped with a prototype small particle side scatter module.

The BD Influx flow cytometer was modified and optimized for detection of submicron-sized particles [10]. In brief, FITC was excited with a 488 nm laser (Sapphire, Coherent 200 mW) and fluorescence was collected through a 530/40 bandpass filter. PE was excited with a 562 nm laser (Jive, Cobolt 150 mW) and fluorescence was collected through a 585/42 bandpass filter. Optical configuration of the forward scatter detector was adapted by mounting a smaller pinhole and an enlarged obscuration bar in order to reduce optical background. This reduced wide-angle FSC (rwFSC) allowed detection of sub-micron particles above the background based on forward scatter [10, 16]. Upon acquisition, all scatter and fluorescence parameters were set to a logarithmic scale. To minimize day to day variations, the BD Influx was standardized at the beginning of each experiment by running 100 and 200 nm yellow-green (505/515) FluoSphere beads (Invitrogen, F8803 and F8848). The instrument was aligned until predefined MFI and scatter intensities where reached with the smallest possible coefficient of variation (CV) for rwFSC, SSC and fluorescence. After optimal alignment, PMT settings required no or minimal day to day adjustment and ensured that each measurement was comparable. MESF beads and NPs were measured with a FSC threshold set at 1.0 while for biological EVs a fluorescence threshold was set at 0.67 by allowing an event rate of 10-20 events/second while running a clean PBS control sample.

When performing quantitative and qualitative analysis of synthetic NPs and biological EVs, preparations were diluted in PBS as indicated. Upon loading on the Influx, the sample was boosted into the flow cytometer until events appeared, after which the system was allowed to stabilize for 30 seconds. Measurements were performed either by a fixed 30 second time or by setting a gate around the spike-in beads and allowing to record a defined number of events in the gate (80 000 events) using BD FACS Sortware 1.01.654 (BD Biosciences).

The CytoFLEX LX was used without any tailor-made modifications in the configuration. Before measurements, the manufacturer recommended startup and QC procedure were run first. All scatter and fluorescence parameters were set to a logarithmic scale. FITC was measured with a 50 mW 488 nm laser and fluorescence was measured through a 525/40 band pass filter at gain 1.0. FITC MESF beads were recorded with an FSC threshold at 1000. Measurements were performed using CytExpert 2.1 (Beckman Coulter).

The SORP BD FACSCelesta^™^ was equipped with a prototype small particle SSC module for improved scatter detection. Before measurement, the recommended CS&T performance check was run to monitor performance on a daily basis and to optimize laser delay. All scatter and fluorescence parameters were set to a logarithmic scale. 100 nm yellow-green (505/515) FluoSphere beads (Invitrogen, F8803) were acquired and used to set optimal fluorescence (FITC detector) PMT-V values. FITC was measured with a 100 mW 488 nm laser through a 530/30 band pass filter. FITC-MESF beads were recorded with an SSC threshold at 200. Measurements were performed using BD FACSDiva^™^ Software v8.0.3 (BD Biosciences).

Further descriptions of each instrument and methods are provided in Data S1 (MIFlowCyt checklist) and Data S2 (MiFlowCyt-EV framework).

### Preparation of FITC-doped silica nanoparticles

Synthetic silica nanoparticles (SiNPs) of 550 nm diameter with six different FITC fluorescence intensities were produced by using a modified method of literature reports [17–19]. Briefly, the amine reactive FITC molecules were covalently linked to the silane coupling agent, (3-aminopropyl)-triethoxylsilane (APTES) in anhydrous ethanol. Monodisperse silica seeds of ~90 nm prepared by using amino acid as the base catalyst [17, 18] were suspended in a solvent mixture containing ethanol, water and ammonia. Then tetraethyl orthosilicate (TEOS) and different volumes of APTES-FITC solutions were added for growing FITC-doped SiNPs by a modified Stöber method [19, 20]. Upon washing three times with anhydrous ethanol, the FITC-doped SiNPs were reacted with TEOS in the solvent mixture to allow growth of a silica layer. The synthesized SiNPs were washed three times with anhydrous ethanol and stocked in anhydrous ethanol. The diameters of SiNPs were measured by transmission electron microscopy.

### Isolation and fluorescent staining of extracellular vesicles for flow cytometric analysis

EV-containing samples were obtained from 4T1 mouse mammary carcinoma cell culture supernatants (ATCC, Manassas, VA) as previously described [16, 21, 22]. EVs were stained with 5-(and-6)-Carboxyfluorescein diacetate succinimidyl ester (CFDA-SE, hereinafter referred as CFSE) (Thermo Fisher, Catalog No. C1157) and separated as described previously [10]. Briefly, 2 μl of the isolated 4T1 EVs (corresponding to a concentration of 1.44 E12 particles/mL as determined by nanoparticle tracking analysis) were mixed with 18 μl PBS/0.1% aggregate-depleted (ad)BSA. For antibody labeling, samples were first resuspended in 15.5 μL PBS/0.1% adBSA and incubated with 0.5 μg of rat anti-mouse CD9-PE (Clone: KMC8, IgG2a, κ lot 7268877, BD Biosciences) or matched isotype antibodies (Rat IgG2a, κ PE-conjugated, lot 8096525, BD Biosciences) for 1h at RT while protected from light. EVs were then stained with 40 μM CFSE in a final volume of 40 μl. The sealed tube was incubated for 2h at 37°C while protected from light. Next, staining was stopped by adding 260 μl PBS/0.1% adBSA. After fluorescent staining, EVs were separated from protein aggregates and free reagents by bottom-up density gradient centrifugation in sucrose for 17.30 h at 192,000 g and 4°C using a SW40 rotor (k-factor 144.5; Beckman Coulter, Fullerton, California, USA). Twelve fractions of 1 mL were then collected from the top of the gradient and respective densities were determined by refractometry using an Atago Illuminator (Japan). For analysis by flow cytometry, EV samples corresponding to a 1.14 g/mL density were diluted 1:20 in PBS prior measurement. MIFlowCyt-EV framework [8] were followed whenever applicable (Data S2).

### Concentration determination by using spike-in beads

EV concentration was normalized using a spiked-in external standard containing 200 nm orange (540/560) fluorescent beads (Invitrogen, F8809). The concentration of the beads was determined by Flow NanoAnalyzer N30 (NanoFCM, Xiamen, China) and stocked at 5.7E10 particles/mL. Beads were diluted 1:10^4^ in PBS and added to the EV samples, mixed and measured on the flow cytometer. Bead count was used to calculate the EV concentration for BD Influx measurements.

## Data analysis

For fluorescence calibration each bead peak population was gated using FlowJo Version 10.5.0 and MFI were obtained for further least square linear regression analysis. Data was handled in Microsoft Excel and figures were prepared using GraphPad Prism version 8.0 (GraphPad Software Inc). The software FCMPASS Version v2.17 was used to generate files with calibrated axis units in the histograms and dot plots shown [23] (Software is available on http://go.cancer.gov/a/y4ZeFtA).

## Data availability

All EV data of our experiments have been submitted to the EV-TRACK knowledgebase (EV-TRACK ID: EV210047) [24]. All flow cytometric data files have been deposited at the Flow Repository (FR-FCM-Z3FJ).

## Funding Statement

This research is supported by the European Union’s Horizon 2020 research and innovation programme under the Marie Skłodowska-Curie grant agreement No 722148 and by the National Natural Science Foundation of China (21934004 and 21627811). E. L. A. is supported by the European Union’s Horizon 2020 research and innovation programme under the Marie Skłodowska-Curie grant agreement No 722148.

## Author Contributions

E.L.A. designed and performed experiments, analyzed data and wrote the manuscript. T.B. performed experiments and gave conceptual advice. L.W. gave technical and conceptual advice. M.M., Y.T. and X.Y. prepared materials and gave technical advice. G.J.A.A. and M.H.M.W. supervised the research, designed (performed) experiments and wrote the manuscript. G.J.A.A. and M.H.M.W. contributed equally as senior author. All authors critically reviewed and edited the manuscript.

## Acknowledgments

The authors would like to thank Prof. An Hendrix (Laboratory of Experimental Cancer Research, Ghent University, Belgium) for the possibility to prepare and analyze M4T1 derived EV in her lab, Dr. Joshua A. Welsh (National Cancer Institute, Bethesda, MD) for helpful discussion and Ludo Monheim (BD Biosciences, Erembodegem, Belgium) for helpful technical support. FITC-MESF and 2 μm PE-MESF beads were kindly provided by BD Biosciences (prepared by Dr. Majid Mehrpouyan) as part of the European Union’s Horizon 2020 research and innovation programme under the Marie Skłodowska-Curie grant agreement No 722148.

## Declaration of interest disclosure

Tina Van Den Broeck and Majid Mehrpouyan are both employees of BD Biosciences, a business unit of Becton, Dickinson and Company. During the course of this study, the Wauben research group, Utrecht University, Faculty of Veterinary Medicine, Department of Biomolecular Health Sciences and BD Biosciences collaborated as a co-joined partner in the European Union’s Horizon 2020 research and innovation programme under the Marie Skłodowska-Curie grant agreement No 722148. Xiaomei Yan declares competing financial interests as a cofounder of NanoFCM Inc., a company committed to commercializing the nano-flow cytometry (nFCM) technology.

## Supporting Information

Additional supporting information can be found online in the corresponding section at the end of the article.

**Data S1.** Author Checklist: MIFlowCyt-Compliant Items.

**Data S2.** MIFlowCyt-EV framework.

**Figure S1.** Comparison of the inclusion of different data points into linear regression analysis by using custom-made and commercial FITC MESF bead sets.

**Figure S2.** Analysis of different MESF bead sets measured on the BD Influx.

**Figure S3.** Analysis of instrument linearity on the BD Influx.

**Figure S4.** Evaluation of measurement variability of 2 μm FITC MESF beads measured on the BD Influx.

**Figure S5.** Gating strategy for synthetic silica NPs.

**Figure S6.** 200 nm fluorescent polystyrene NPs.

**Figure S7.** Gating strategy for external spiked-in beads and biological EV samples.

## Notes

### Summary of Updates

Main text has been updated Figures have been revised and updated

