## Supplementary material for "Intrinsic variability of fluorescence calibrators impacts the assignment of MESF or ERF values to nanoparticles and extracellular vesicles by flow cytometry": Supplmental material

**Data S1. Author Checklist: MIFlowCyt-Compliant Items.**

| Requirement | Please Include Requested Information |
|---|---|
| 1.1. Purpose | Evaluate the advised use of MESF bead-based calibration for the assignment of absolute fluorescent values to nanoparticles and extracellular vesicles. |
| 1.2. Keywords | flow cytometry, fluorescence, calibration, standardization, MESF, extracellular vesicles, nanoparticles |
| 1.3. Experiment variables | MESF bead, flow cytometer |
| 1.4. Organization name and address | Department of Biomolecular Health Sciences, Faculty of Veterinary Medicine,<br>Yalelaan 2 (Nieuw Gildestein Room 203), University of Utrecht<br>PO Box 80176,<br>3508 TD Utrecht,<br>The Netherlands |
| 1.5. Primary contact name and email address | Estefanía Lozano-Andrés<br>Marca H. M. Wauben |
| 1.6. Date or time period of | January 2019 until October 2021 |

| | |
|---|---|
| experiment | |
| 1.7. Conclusions | By testing various MESF bead sets we found differences in the slopes of the regression lines consistent between different calibrator bead sets and not dependent on the flow cytometer platform used. These differences are caused by uncertainties in the assignment of MESF to the calibrators and are exaggerated during extrapolation through linear regression into the dimmer fluorescent area. |
| 1.8. Quality control measures | 100 nm polystyrene beads, 200 nm polystyrene beads, QC samples as provided by manufacturers. |
| 2.1.1.1. Sample description | Fluorescent 100 nm polystyrene beads, Fluorescent 200 nm polystyrene beads, two different sets of FITC MESF beads (6 µm and 2 µm), two different sets of PE MESF beads (6 µm and 2 µm), 550 nm Silica Nanoparticles FITC-labeled, 200 nm fluorescent polystyrene beads for concentration determination, extracellular vesicles (EVs) isolated from 4T1 cell culture supernatant. |
| 2.1.1.2. Biological sample source description | Murine mammary carcinoma cell line 4T1 (ATCC, Manassas, VA). |
| 2.1.1.3. Biological sample source organism description | Mouse |
| 2.1.2.2. Environmental sample location | NA |
| 2.3. Sample treatment description | EV samples were succumbed to differential ultracentrifugation, ultrafiltration and density gradient floatation. |
| 2.4. Fluorescence reagent(s) description | EV sample were stained with with 5-(and-6)-Carboxyfluorescein Diacetate Succinimidyl Ester (CFDA-SE, hereinafter referred as CFSE) (ThermoFisher, catalog number C1157) and labeled with 0.5 µg of Rat anti-mouse CD9-PE (Clone: KMC8, IgG2a, κ, Lot. no. 7268877, BD Biosciences) or matched Isotype antibodies (Rat IgG2a, κ, PE-conjugated, Lot. no. 8096525, BD Biosciences). |
| 3.1. Instrument manufacturer | BD Biosciences, Beckman Coulter. |
| 3.2. Instrument model | BD Influx™, BC CytoFLEX LX™ and SORP BD FACSCelesta™ |
| 3.3. Instrument configuration and settings | The BD Influx™ was optimized for detection of sub-micron sized particles as described previously in Van der Vlist *et al.* – Nature Protocols 2012. Based in that configuration various combinations of pinholes and obscuration bars were tested as described in the manuscript. Prior to each measurement, 100 nm yellow-green (505/515) FluoSphere beads (Invitrogen, F8803) were used to set optimal values. FITC was measured with a |

| | |
|---|---|
| | 200 mW 488nm laser (Sapphire, Coherent) placed in front of the first pinhole and through a PMT 530/40 band pass filter. PE was measured with a 150 mW 561nm laser (Jive, Cobolt) placed in front of the third pinhole and through a PMT 585/42 band pass filter. FITC MESF and PE MESF beads were recorded with an FSC threshold at 1.00. Synthetic Nanoparticles and Biological EV samples were recorded with a FL threshold at 0.67.<br>CytoFLEX LX™ was used without any tailor-made modifications. Standard startup and QC procedure were run prior measurements as recommended by the manufacturer. All scatter and fluorescence parameters were set to a logarithmic scale. FITC was measured with a 50 mW 488 nm laser and fluorescence was measured through an APD 525/40 band pass filter. Samples were recorded with a FSC threshold at 1000.<br>The SORP BD FACSCelesta™ was equipped with a small particle SSC module for improved scatter detection. Before measurement, the standard setup and QC procedures were performed. All scatter and fluorescence parameters were set to a logarithmic scale. 100 nm yellow-green (505/515) FluoSphere beads (Invitrogen, F8803) were acquired and used to set optimal small particle-SSC and fluorescence (FITC detector) PMT-V values. FITC was measured with a 100 mW 488 nm laser through a PMT 530/30 band pass filter. FITC-MESF beads were recorded with an SSC threshold at 200. |
| 4.1. List-mode data files | The flow repository ID for peer-review process: FR-FCM-Z3FJ |
| 4.2. Compensation description | No compensation was applied. |
| 4.3. Data transformation | No data transformation was applied. |
| 4.4.1. Gate description | Gates were defined by using FlowJo Version 10.5.0.<br>Briefly, for MESF bead gating on the BD Influx™ singlets were gated based on the parameters trigger pulse width Vs SSC, then a histogram displaying fluorescence was plotted to gate the individual peak populations, SiNPs and EVs gating strategy can be found in the Supporting Information of this manuscript (Figure S3 and S5, respectively). For MESF bead gating on the CytoFLEX LX™ singlets were gated based on SSC Vs FSC, then a histogram displaying fluorescence was plotted to gate the individual peak populations. For MESF bead gating on the SORP BD FACSCelesta™ singlets were gated based on SSC Vs FSC, then a histogram displaying fluorescence was plotted to gate the individual peak populations. |
| 4.4.2. Gate statistics | Plots show the MFI (Median Fluorescence Intensity) for each population on the described parameter. See figures of the manuscript and results section for further details. |
| 4.4.3. Gate boundaries | Gates boundaries were set according to each population. See additional Supporting Information. |

**Data S2. MIFlowCyt-EV framework.**

| | |
|---|---|
| 1.1 Preanalytical variables conforming to MISEV guidelines | Yes, all relevant data has been submitted to EV-TRACK for transparent reporting and centralizing knowledge in extracellular vesicle research (EV-TRACK ID: EV210047). |

| | |
|---|---|
| 1.2 Experimental design according to MIFlowCyt guidelines | Yes, MIFlowCyt checklist can be found as part of the supporting information of this manuscript (Data S1). |
| 2.1 Sample staining details | Yes, described in Materials and Methods. |
| 2.2 Sample washing details | Yes, described in Materials and Methods. |
| 2.3 Sample dilution details | Yes, described in Materials and Methods. |
| 3.1 Buffer-only controls | Yes, relevant buffer controls were measured. |
| 3.2 Buffer with reagent controls | Yes, see Figure S5. |
| 3.3 Unstained controls | Yes, see Figure 3. |
| 3.4 Isotype controls | Yes, see Figure 3. |
| 3.5 Single-stained controls | N/A |
| 3.6 Procedural controls | Yes, see Figure S5b. |
| 3.7 Serial dilutions | Yes, serial dilutions were performed in previous characterization experiments to determine the ideal dilution used in this study. |
| 3.8 Detergent-treated controls | Yes, sensitivity to triton X-100 for this EV sample was previously determined. |
| 4.1 Trigger channel(s) and threshold(s) | Yes, all relevant details can be found in Materials and Methods. |
| 4.2 Flow rate / volumetric quantification | Yes, low flow rate was kept constant and is described in Materials and Methods. |
| 4.3 Fluorescence calibration | Yes |
| 4.4 Scatter calibration | N/A |
| 5.1 EV diameter/surface area/volume approximation | N/A |
| 5.2 EV refractive index approximation | N/A |
| 5.3 EV epitope number approximation | N/A |
| 6.1 Completion of MIFlowCyt checklist | Yes, see Data S1 |
| 6.2 Calibrated channel detection range | BD Influx |
| 6.3 EV number/concentration | Yes, see Supporting Information Figure S5. |
| 6.4 EV brightness | Yes, reported in ERF or MESF, see Figure 3. |
| 7.1 Sharing of data to a public repository | Yes, all experimental details can be found in EV-TRACK. Data files have been submitted to http://flowrepository.org under FR-FCM-Z3FJ and are available upon request |

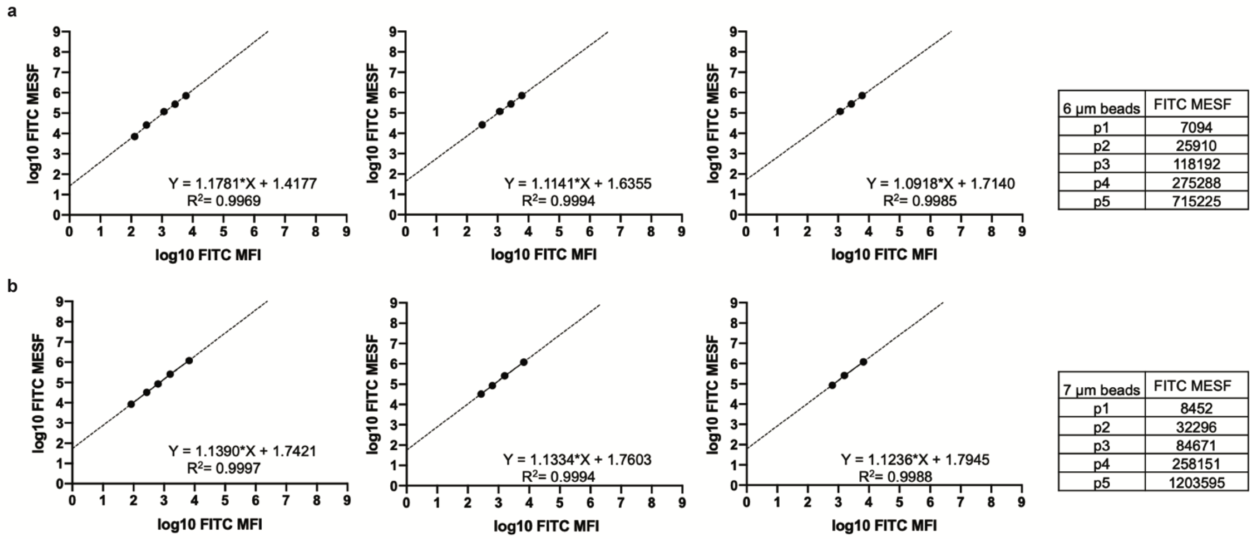


**Figure S1. Comparison of the inclusion of different data points into linear regression analysis by using custom-made and commercial FITC MESF bead sets. (a)** Least square linear regression analysis of custom-made 6 µm FITC MESF beads and **(b)** 7 µm Quantum$^{TM}$ FITC-5 MESF beads measured on the BD Influx. Provided FITC MESF for both sets (indicated in the figure) and measured FITC MFI values were transformed to log and plotted in a log-log fashion. Graphs including either five, four or three fluorescent populations (from left to right).

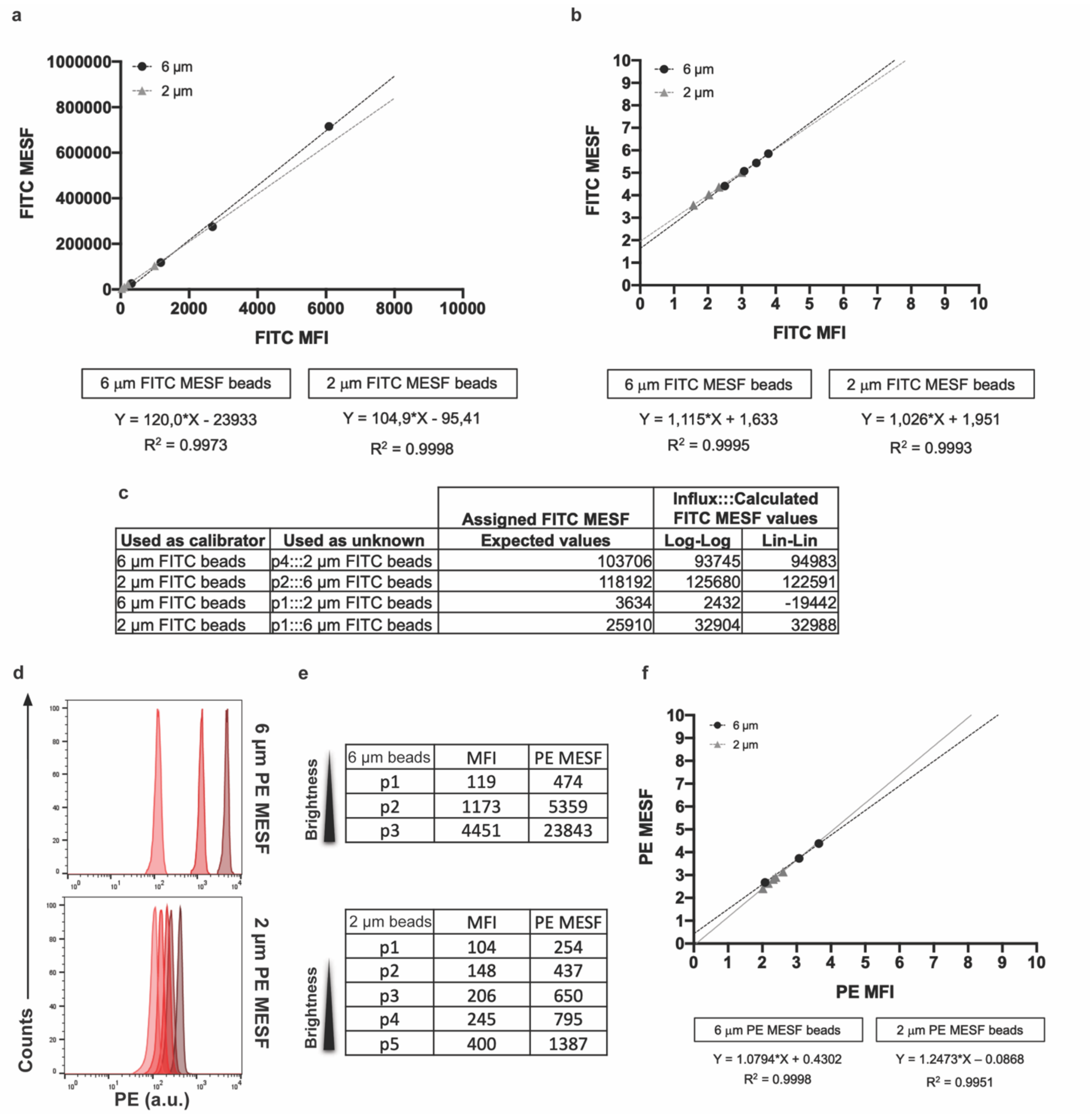


| Used as calibrator | Used as unknown | Assigned FITC MESF Expected values | Influx:::Calculated FITC MESF values Log-Log | Influx:::Calculated FITC MESF values Lin-Lin |
|---|---|---|---|---|
| 6 µm FITC beads | p4:::2 µm FITC beads | 103706 | 93745 | 94983 |
| 2 µm FITC beads | p2:::6 µm FITC beads | 118192 | 125680 | 122591 |
| 6 µm FITC beads | p1:::2 µm FITC beads | 3634 | 2432 | -19442 |
| 2 µm FITC beads | p1:::6 µm FITC beads | 25910 | 32904 | 32988 |

| 6 µm beads | MFI | PE MESF |
|---|---|---|
| p1 | 119 | 474 |
| p2 | 1173 | 5359 |
| p3 | 4451 | 23843 |

| 2 µm beads | MFI | PE MESF |
|---|---|---|
| p1 | 104 | 254 |
| p2 | 148 | 437 |
| p3 | 206 | 650 |
| p4 | 245 | 795 |
| p5 | 400 | 1387 |

**Figure S2. Analysis of different MESF bead sets measured on the BD Influx. (a)** Least square linear regression analysis of 6 µm (black circles) and 2 µm (grey triangles) FITC MESF beads by using linear plotting or **(b)** log-transformed plotting. **(c)** Table indicating the assigned FITC MESF values for each population and the calculated FITC MESF values for each sample. **(d)** Histogram overlays of PE fluorescent intensity peaks derived from the 6 µm or the 2 µm PE MESF beads (arbitrary units). **(e)** Table showing the median intensity fluorescence (MFI) statistic derived from each of the fluorescent intensity peaks from dimmer to brighter expressed in arbitrary units. **(f)** Least square linear regression analysis of 6 µm (black circles) and 2 µm (grey triangles) PE MESF beads. Provided PE MESF and measured PE MFI values were transformed to log and plotted in a log-log fashion.

**a**

| | 488 nm laser | | |
|---|---|---|---|
| | Geo Mean (FITC) | | |
| PMT1 voltage (530/30) | p2 | p6 | p2-p6 |
| 41.85 | 5719 | 104 | 5615 |
| 38.85 | 3287 | 57.2 | 3229.8 |
| 35.85 | 1923 | 31.6 | 1891.4 |
| 32.85 | 963 | 15.7 | 947.3 |

**b**

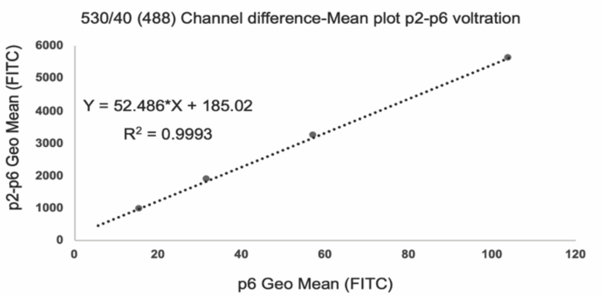


**c**

| | 561 nm laser | | |
|---|---|---|---|
| | Geo Mean (PE) | | |
| PMT11 voltage (585/40) | p5 | p6 | p5-p6 |
| 71.1 | 300 | 106 | 194 |
| 68.1 | 222 | 71.1 | 150.9 |
| 65.1 | 158 | 48.8 | 109.2 |
| 62.1 | 108 | 31.1 | 76.9 |

**d**

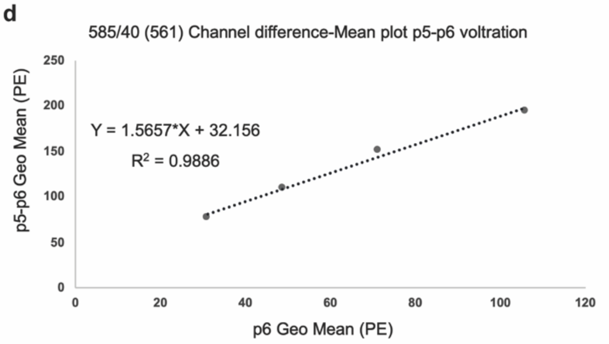


**Figure S3. Analysis of instrument linearity on the BD Influx. (a)** Table indicating the PMT1 voltage and the FITC MFI values for either population p2 or p6 from the beads. **(b)** Linear regression analysis showing the MFI difference between population p2 and p6 relative to the MFI from population p6 and the respective equation. **(c)** Table indicating the PMT11 voltage and the PE MFI values for either population p5 or p6 from the beads. **(d)** Linear regression analysis showing the MFI difference between population p5 and p6 relative to the MFI from population p6 and the respective equation.

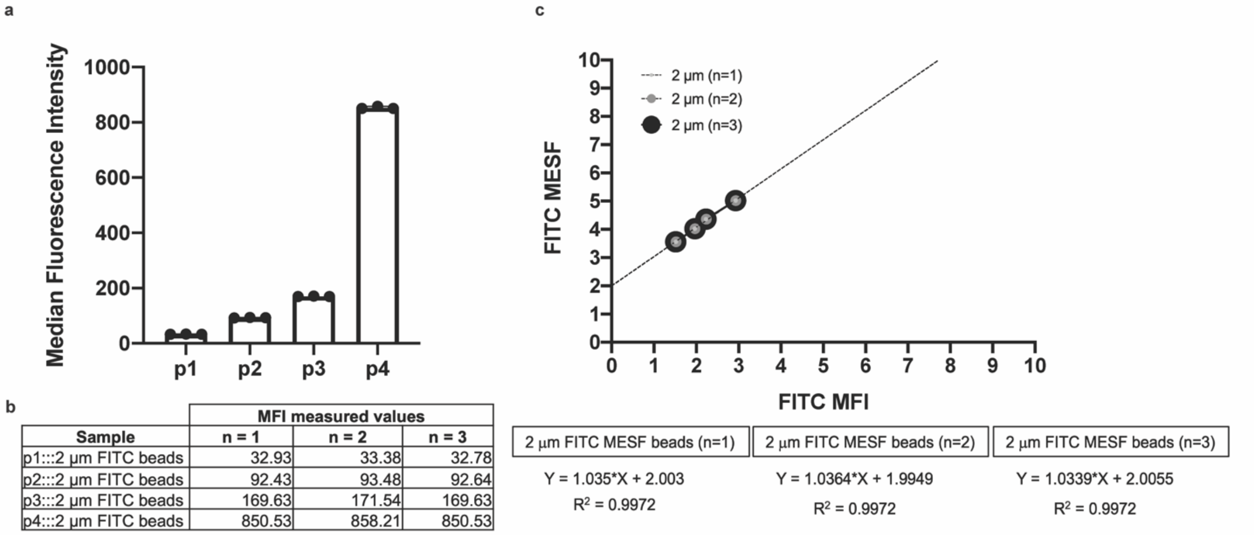


| | MFI measured values | | |
|---|---|---|---|
| Sample | n = 1 | n = 2 | n = 3 |
| p1:::2 µm FITC beads | 32.93 | 33.38 | 32.78 |
| p2:::2 µm FITC beads | 92.43 | 93.48 | 92.64 |
| p3:::2 µm FITC beads | 169.63 | 171.54 | 169.63 |
| p4:::2 µm FITC beads | 850.53 | 858.21 | 850.53 |

**Figure S4. Evaluation of measurement variability of 2 µm FITC MESF beads measured on the BD Influx. (a)** Bar graph displaying the MFI values for the four fluorescent bead populations present in the 2 µm FITC MESF bead set from three independent measurements. **(b)** Table indicating the FITC MFI values for each population from three independent measurements. **(c)** Least square linear regression analysis showing three independent measurements of the 2 µm FITC MESF beads by using log-transformed plotting and their respective equations.

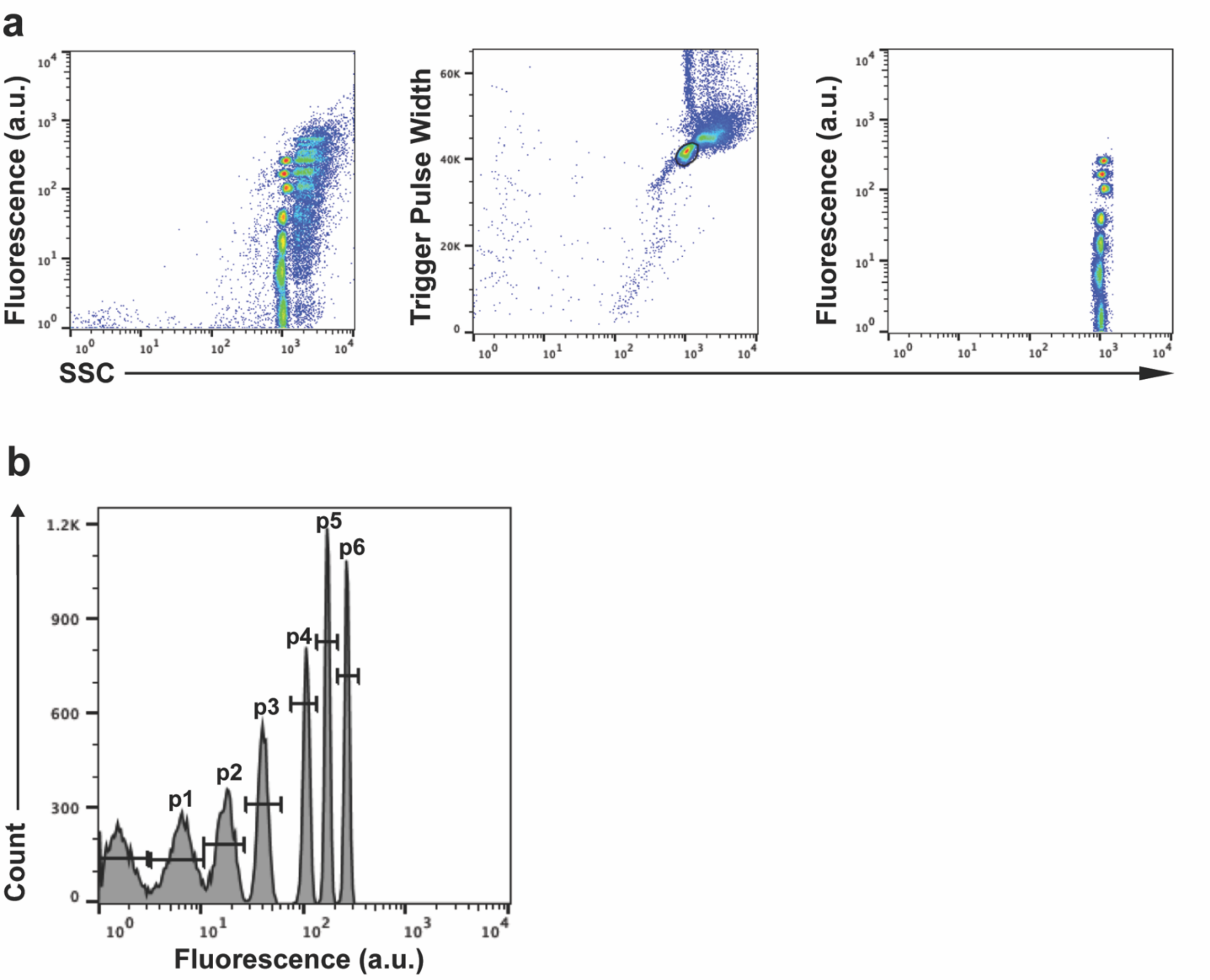


**Figure S5. Gating strategy for synthetic silica NPs. (a)** Dot plots showing fluorescence in arbitrary units vs SSC from 550 nm silica NPs. Ungated events are displayed in a fluorescence vs SSC dot plot (left panel). A gate was placed around the singlet population based on trigger pulse width vs SSC (mid panel). Singlets displaying six FITC intensities and a blank NP population are shown in the Fluorescence Vs SSC (right panel). **(b)** Histogram showing the gated 6 fluorescence intensities (p1-p6) and the blank NP population.

**a**

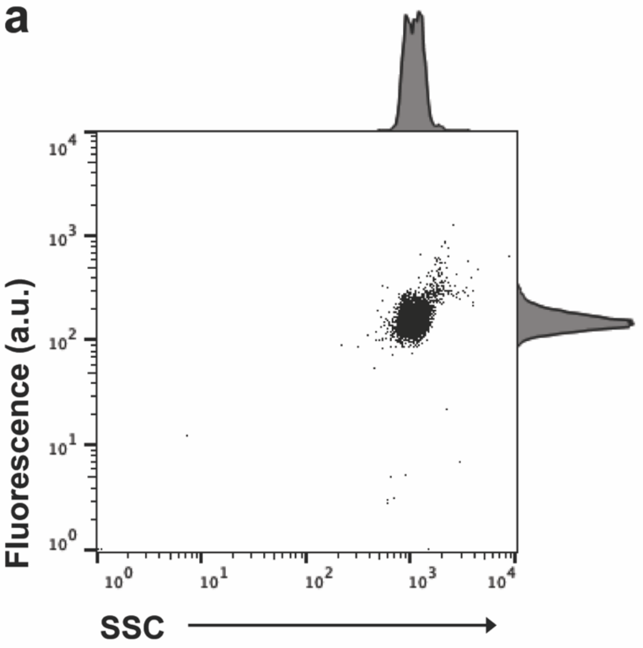


**b**

| | | Calculated ERF values | | |
|---|---|---|---|---|
| | | Used as calibrator | | |
| Unknown | MFI | 6 µm PE beads | 2 µm PE beads | % variation |
| 200 nm NPs | 154 | 619 | 438 | 41.3 |

**Figure S6. 200 nm fluorescent polystyrene NPs. (a)** Dot plots showing fluorescence in arbitrary units (a.u.) in the PE channel vs SSC from a sample containing 200 nm broad spectrum fluorescent polystyrene synthetic NPs. **(b)** Table indicating the ERF values for the 200 nm fluorescent NPs calculated either using 6 or 2 µm PE MESF beads.

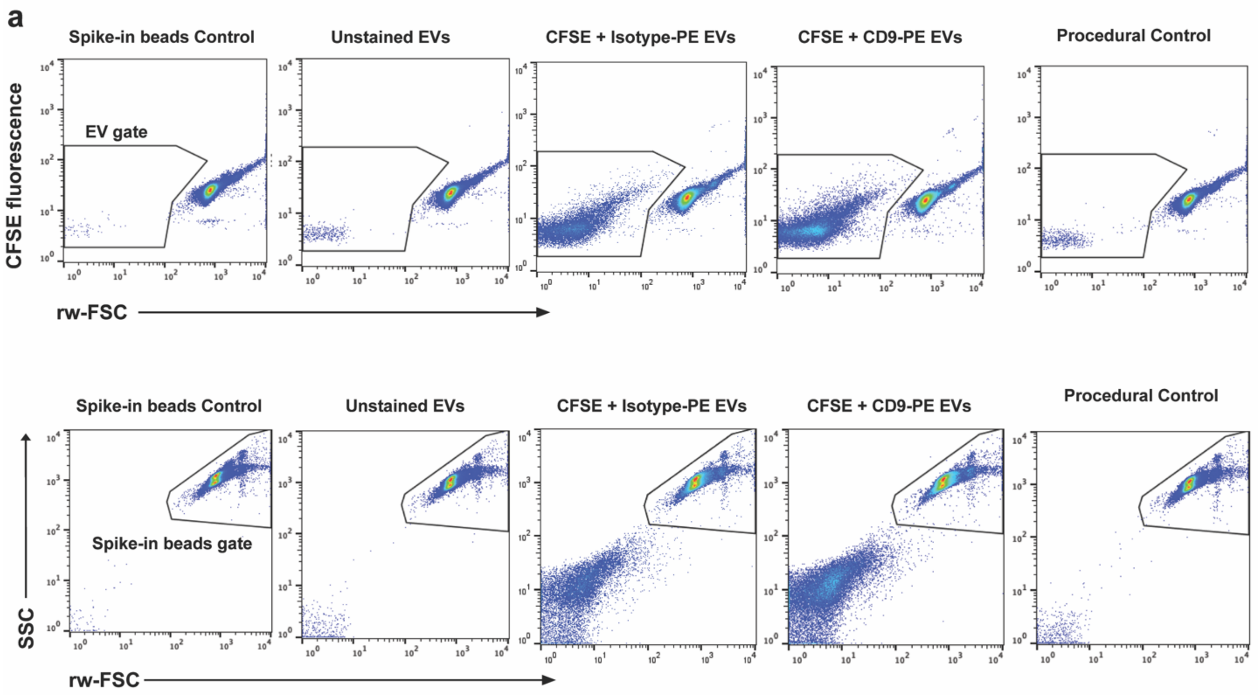


**Figure S7. Gating strategy for external spiked-in beads and biological EV samples. (a)** Dot plots showing fluorescence in arbitrary units (a.u.) vs rw-FSC or SSC vs rw-FSC from a spike-in beads control, an unstained EVs control, an isotype EVs control, a CFSE-stained and CD9-PE labeled EV sample and a procedural control. The concentration of the CFSE stained and CD9-PE labeled EV sample was calculated using the spike-in beads (previously described in Material and Methods) and estimated to be $2.34 \times 10^7$ particles/mL.
